# Exploring the association between Body Mass Index, Sex and Gene Expression in human colorectal epithelium

**DOI:** 10.1101/2022.11.21.515057

**Authors:** L. Lemler, K. Donnelly, I. P. M. Tomlinson, M. Timofeeva, E. Theodoratou, C. Fernández Rozadilla, J. Fernandez-Tajes, Graeme Grimes, Susan M. Farrington, M. G. Dunlop

**Author notes:** **Correspondence:** Professor MG Dunlop. Institute of Genetics and Cancer, University of Edinburgh and MRC Human Genetics Unit, Western General Hospital Edinburgh, Crewe Road, Edinburgh, EH4 2XU, United Kingdom. E.

## Abstract

**Introduction:** Colorectal cancer (CRC) is the second most common cause of cancer death globally. Genome-wide association studies have established that cancer risk mediated through common genetic variants can be linked to variation in gene expression. Since obesity and male sex impart substantially elevated CRC risk, we studied transcriptional profiles of normal colorectal mucosa using RNA sequencing to better understand the relationship of these risk factors with gene expression levels.

**Methods:** Normal colorectal mucosa was sampled from 365 participants (208 males, 157 females) either during surgery (n=103) or through endoscopic biopsy (n=262) from cancer patients and patients with other unrelated conditions. In total, 238 samples were used for our discovery dataset and 380 samples were obtained for the validation of our findings. The transcription analysis was done using paired-end total RNA sequencing. Data processing and gene filtering followed the Genotype-Tissue Expression (GTEx) Project pipeline v8. Differential Expression Analysis (DEA) was performed on normalised counts to evaluate effects of sex and body mass index on the total gene expression, as well as possible confounding effects of cancer presence on the gene expression in normal colorectal tissue.

**Results:** Following filtering, there were 15,465 genes available for analysis. DEA identified two genes that were significantly associated with sex and five associated with body mass index. However, whilst these nominal signals are of interest, none of the genes associated with sex remained significant in a replication dataset. Due to the missing BMI information, replication of DEA by BMI was not possible.

**Conclusion:** We found no systematic differences in gene expression in normal colorectal epithelium between males and females, nor did we find a strong association between gene expression and BMI. Although sample size may limit our analysis, the results suggest no or limited confounding effects of BMI and sex on gene expression in normal colorectal mucosa samples.

## INTRODUCTION

Colorectal Cancer (CRC) has the third highest incidence among different types of cancer globally and is the second most common cause of cancer related death worldwide (accounting for 9.2% of all cancer deaths across the globe annually) (Bray *et al*., 2018).

Sex and body mass index (BMI) substantially influence the risk of developing CRC with pooled relative risk (RR) of overweight and obese men at 1.37 (95% CI: 1.21-1.56) and 1.07 for women (95% CI: 0.97-1.18) when compared to the same BMI group (Dai, Xu and Niu, 2007). According to a recent World Health Organisation report, more than 1.9 billion adults are overweight or obese *(World Cancer Report: Cancer Research for Cancer Prevention*, 2022). 11% of CRC cases have been associated to overweight and obesity in Europe, which makes it one of the most common obesity-related cancers (Tarasiuk, Mosińska and Fichna, 2018) (Bardou, Barkun and Martel, 2013). While obesity is a risk factor for both sexes, this effect seems stronger among men (Kim and Giovannucci, 2017). Despite this supportive evidence, it is still not clear why obese individuals as well as males have a significantly higher CRC risk.

High-throughput technologies have enabled the analysis of gene expression in large cohorts and opened new opportunities for investigation. Recent progress in the understanding of gene expression and its relationship to genetics have shown that inherited cancer risk can be linked to variation in gene expression (Loo, Lemire and Le Marchand, 2017).

An analysis of healthy individuals further suggests that differences in gene expression in whole blood are associated with BMI (Joseph *et al*., 2019). Moreover, a limited number of recent studies have shown sex differences in the transcriptome of normal colorectal mucosa (Hases *et al*., 2021) (Oliva *et al*., 2020) (Naqvi *et al*., 2019) (Melé *et al*., 2015) (Gershoni and Pietrokovski, 2017). However, the majority of these studies use the same dataset (GTEx) or have very limited sample sizes (*GTEx Portal*, 2022).

Previous studies of gene expression have mainly concentrated on tumour tissue and the comparison between tumour tissue and adjacent normal tissue. Variation in gene expression in normal tissue has been less studied, but it could play an important role in understanding the differences among individuals (Huang, Stern and Zhao, 2016). Huang (2016) even suggests that transcriptional profiles of normal tissue are more informative as predictors of patient survival than tumour tissue. A previous publication by our group analysed the gene expression differences in site of normal mucosa of 481 subjects (Vaughan-Shaw *et al*., 2021). 1261 differentially expressed genes were identified with the top hit showing a fold-difference of 10. Moreover, larger differences in gene expression between sex in normal mucosa compared to CRC tissue have been reported (Hases *et al*., 2021).

Here we present an exploratory RNA Sequencing analysis of normal colorectal mucosa. The aim of this study is to describe the gene expression differences associated with sex and BMI in normal colorectal mucosa, which could be crucial for understanding obesity and sex specific differences in CRC risk. It could have further implications on screening, prevention and treatment of CRC.

## MATERIALS AND METHODS

### Study Subjects and Normal Mucosa Samples

Colorectal normal mucosa samples of a subset of individuals of the Study of Colorectal Cancer in Scotland (SOCCS) (Fernandez-Rozadilla *et al*., 2022) were included for gene expression analysis. The samples were collected at the Western General Hospital, Edinburgh between April 2010 and March 2019. The samples were taken from a single site during surgery (stripped mucosa) or through mucosal rectal biopsy from cancer patients and patients with minor ailments or investigations unrelated to colorectal cancer. However, as shown in a previous paper, no difference has been observed between both groups (P.G. Vaughan-Shaw 2022). Sigmoid and Descending Colon might be overlapping in the SOCCS because the borderline of the location of these samples was hard to define. The basic characteristics of both studies are presented in **Table 1**. Overall, 238 samples were collected, containing 104 samples from females and 134 samples from males. The median age of the study population was 64 years old (overall range 24-89). Most of the individuals were of Northern European (Scottish) descent (with one Chinese, two African and two Middle Eastern ethnicities). PCA plots based on genotyping array data showed these ethnicities accordingly. Further samples were collected from the Intermediate Phenotype Study (INTERMPHEN). Samples were obtained in Oxford between November 2012 and March 2014, and corresponded to 127 individuals of white UK origin (74 males and 53 females) that underwent colonoscopy. From each of these, three samples from the rectum, sigmoid colon and caecum were collected. These samples had a within-patient correlation of gene expression of 0.34. Indications for colonoscopy were CRC and non-CRC related, however none of the patients presented with CRC. The median age was 61 years (overall range 21-80). Discovery and replication datasets had similar proportion of women (SOCCS: 43.7%; INTERMPHEN: 41.7%) and were similar in age, but SOCCS was overrepresented by distal samples.

**TABLE 1.** Characteristics of normal colorectal epithelium samples and participants.

| Characteristics |  | SOCCS n=238 | INTERMPHEN n=127 with<br>3 sample sites for each<br>participant |
| --- | --- | --- | --- |
| Sex | Male | 134 | 74 |
|  | Female | 104 | 53 |
| Age | Median | 64 | 61 |
|  | Range | 24-89 | 21-80 |
|  | Male Range | 24-86 | 23-80 |
|  | Female Range | 24-89 | 21 -75 |
| BMI (kg/m2) | Median | 26.9 | No BMI available. |
|  | Range | 14.7-48.9 |  |
|  | Low (<24.48) | 78 (32.77%) |  |
|  | Medium (24.48-29.34) | 79 (33.19%) |  |
|  | High (>29.34) | 81 (34.03%) |  |
| Cancer Status | None | 109 | 69 |
|  | Adenoma | 10 | 58 |
|  | PMH<sup>1</sup> CRC | 18 |  |
|  | CRC | 100 |  |
|  | Small Bowel Tumour | 1 |  |
| Type of Tissue | Endoscopic Biopsy | 135 | 380 |
|  | Stripped Mucosa | 103 |  |
| Sample Site | **Proximal** | **33** | **127** |
|  | Transverse Colon | 29 |  |
|  | Caecum | 4 | 127 |
|  | **Distal** | **205** | **253** |
|  | Rectum | 144 | 126<sup>2</sup> |
|  | Sigmoid Colon | 12 | 127 |
|  | Descending Colon | 49 |  |

1 Past medical history | 2 Only one sample for rectum

BMI data was only available for SOCCS. Females showed a lower median BMI of 25.5 (range, 14.7-45.7), while the median BMI of males was 27.7 (range, 18.3-48.9). In order to ensure similar grouping of individuals in BMI groups we decided to use tertiles for the BMI groups instead of following the traditional grouping of underweight/normal/overweight/obese 1/obese 2.

All participants of the SOCCS provided informed written consent, and research was approved by local research ethics committees (13/SS/0248) and National Health Service management (2014/0058) for the SOCCS. Ethical approval for the INTERMPHEN study was obtained from the Oxfordshire Research Ethics Committee A and all participants provided informed written consent.

### RNA Extraction

For SOCCS RNA extraction, library preparation and RNA sequencing analysis was performed in three batches. The normal colorectal mucosa samples were surgically stripped from submucosa or taken via rectal biopsy. All samples were immersed in stabilisation solution RNAlater (Invitrogen). A detailed description of the processing has been previously published (Vaughan-Shaw *et al*., 2021).

For the INTERMPHEN study each biopsy was also submerged in a preservation medium containing Dulbecco’s Modified Eagle Medium (DMEM) and RNAlater (Life Technologies) and further treated using standard methodologies as specified previously (Fernandez-Rozadilla *et al*., 2018).

### RNA Sequencing and Data Processing

The RNA sequencing was performed on an Illumina HiSeq 4000 platform (Illumina, San Diego, CA) on 150bp, 75bp and 75bp paired-end reads for SOCCS and 75bp paired-end reads for INTERMPHEN, in accordance with the manufacturer’s instructions.

The data was analysed according to the GTEx Project pipeline v8 (‘Analysis pipelines for the GTEx Consortium and TOPMed’, 2022) **(Figure 1)**. The STAR Alignment (Version 2.5.3a) (Dobin *et al*., 2013) was used to compare the reads with the human reference genome (GRCh38) and to determine where the sequencing reads lie within a transcriptome. Afterwards, the produced bam files were checked and marked for duplicates. Counts were generated on transcript-level using RNA-Sequencing by Expectation-Maximization (RSEM) (Version 1.3.0) (Li and Dewey, 2011) as well as on gene-level using RNA SeQC (Version 2.0.0) (DeLuca *et al*., 2012). GENCODE version v26 was used *(GENCODE - Human Release 26*, 2020). The sequencing generated a median 35 million reads per sample for the SOCCS dataset and 26 million reads per sample for the INTERMPHEN.

**FIGURE 1.**
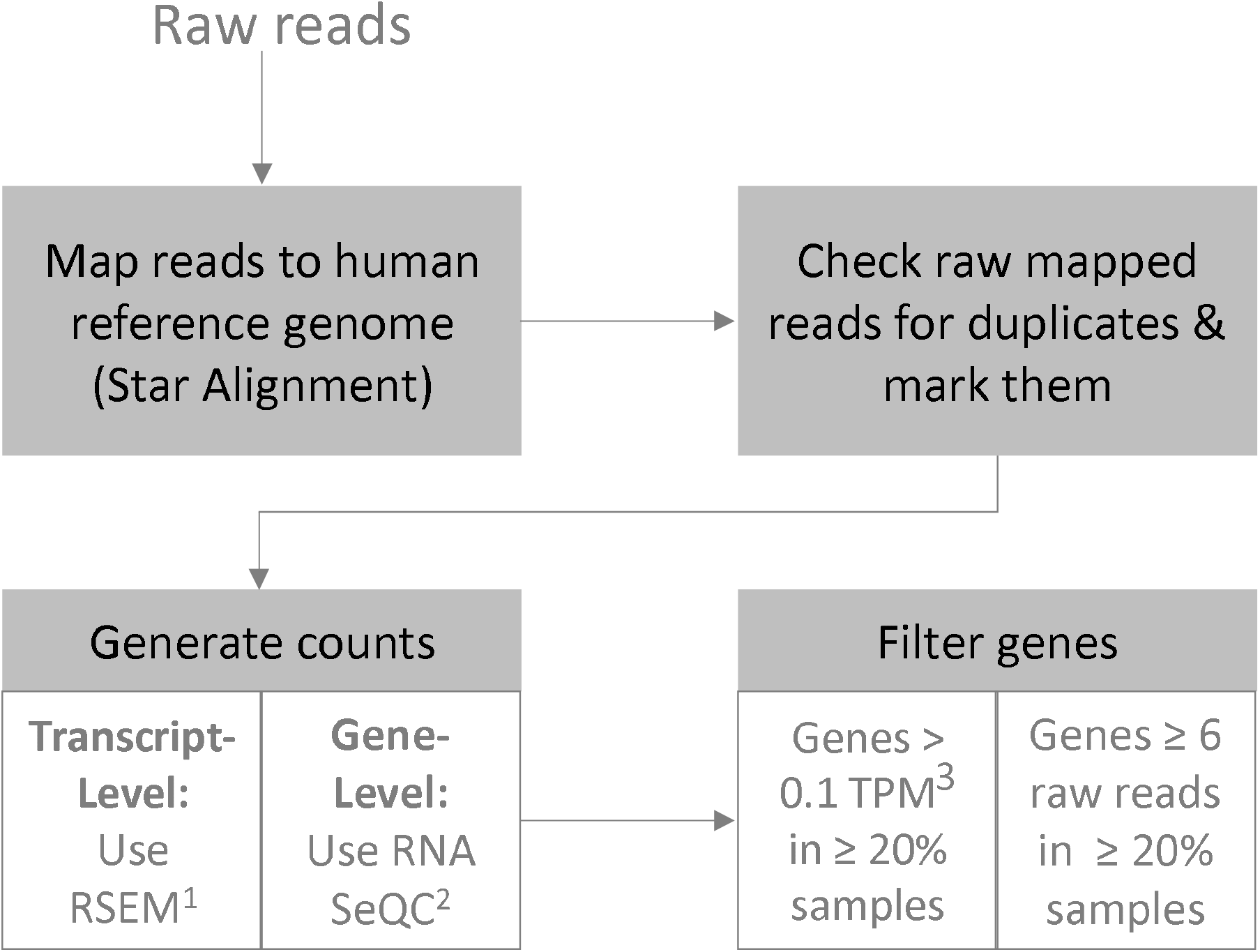
Workflow of RNA Data Processing: Mapping reads, checking raw reads for duplicates, generating counts and filtering the genes. 1 Accurate transcript quantification from RNA-Seq data with or without a reference genome | 2 RNA-seq metrics for quality control and process optimization | 3 Transcript per Million

Gene filtering was performed also according to the GTEx v8 pipeline. Briefly, we kept genes with a transcript per million (TPM) count > 0.1 in at least 20% of the samples and genes with at least 6 raw reads in 20% of the samples. After applying these filters, the SOCCS dataset was reduced from 58,219 to 15,465 genes, whereas INTERMPHEN dropped from 58,219 to 13,771 genes.

The DESeq2 function on raw counts was used to normalise the SOCCS data by taking the natural logarithm of all values as well as calculating the median of the ratios for each sample (Love, Huber and Anders, 2014). The raw INTERMPHEN counts were converted to log2-counts-per-million (logCPM) and the mean-variance relationship was modelled with an empirical Bayes prior trend as suggested in established workflows (Gordon K. Smyth *et al*., 2021).

The analysis was enhanced by filtering the dataset by previously published CRC risk candidates **(Supplementary Table 1**) (Law et al., 2019) as well as 155 effector genes linked to CRC risk (**Supplementary Table 3**) (Fernandez-Rozadilla *et al*., 2022).

## Statistical Methods

### Differential Expression Analysis

The models for the DEA were built using the DESeq2 Package (Version 1.28.1) (Love *et al*., 2022) in R (Version 4.0.2) (*R: The R Project for Statistical Computing*, 2022) and following the workflow specified by the developers (Love, Anders and Huber 2021).

For the INTERMPHEN dataset, linear models were built using the lmfit function of the Limma Package (Version 3.46.0) (Ritchie *et al*., 2015) in R (Version 4.0.2). Since the data included multiple samples from the same patient we utilised the built-in function “duplicateCorrelation” for analysing repeated measures (*dupcor: Correlation Between Duplicates in limma: Linear Models for Microarray Data*, 2022).

The models contained the following factor variables: the variable of interest (sex, BMI, non-cancer vs. cancer, non-cancer vs. neoplasia, non-cancer vs. adenoma) as well as the covariates batch, age, sample site and, where relevant sex, BMI and cancer status to control for potential effects (for Principal-Component-Analysis see **Supplementary Material**). For the stratified analysis for BMI by sex the variable sex was removed. Results were adjusted for multiple testing using the Benjamini-Hochberg procedure for false discovery rate (FDR) (Benjamini and Hochberg, 1995). Differentially expressed genes were defined by an absolute log2 fold-change > 1.2 and a Benjamini-Hochberg– adjusted p value < 0.05 was considered statistically significant (Michael I. Love, Vladislav Kim and Wolfgang Huber, 2019). Significant results on the sex chromosomes were excluded.

## RESULTS

### Association between Sex and Gene Expression

DEA by sex identified several genes on the sex chromosomes with a significant adjusted p-value below 0.05 and log2 fold-change threshold above 1.2, which were excluded. Genes *DUOXA2* and *IGHV2-70D* were the only significant genes not located on X or Y chromosomes (**Table 2**). The gene expression of both genes was higher in females than males.

**TABLE 2.** Results of DEA **male vs. female**. Models have been adjusted for batch, sample site, BMI, age and cancer status. Included samples from SOCCS Study. Included genes with adjusted p-value below 0.05 and log2 fold-change threshold above 1.2.

| Study | Ensembl Gene ID | HGNC<sup>1</sup><br>Symbol | Chromosome | Log2Fold Change | Padj<sup>2</sup> | Pvalue | baseMean<sup>3</sup> | lfcSE<sup>4</sup> |
| --- | --- | --- | --- | --- | --- | --- | --- | --- |
| SOCCS | ENSG00000140274 | DUOXA2 | 15 | -1.22 | 0.01 | 8.25E-05 | 148.29 | 0.31 |
|  | ENSG00000211974 | IGHV2-70D | 14 | -1.21 | 0.01 | 3.93E-05 | 63.24 | 0.29 |
| INTERMPHEN | ENSG00000140274 | DUOXA2 | 15 | 0.17 | 1 | 0.47 | 5.07 | 0.24 |
|  | ENSG00000211974 | IGHV2-70D | 14 | 0.18 | 1 | 0.43 | 4.91 | 0.22 |

1 HUGO Gene Nomenclature Committee | 2 Benjamini-Hochberg – adjusted p value | 3 Average expression of the normalized count values | 4 Standard error estimate for the log2 fold-change estimate

Neither of these genes was replicated in the INTERMPHEN dataset (**Table 2**). We further investigated if any of the known CRC risk candidate genes showed evidence of differential expression by sex. We extracted 46 candidate target genes from the previously published GWAS on CRC risk (**Supplementary Table 1**) (Law et al., 2019). None of the candidate genes showed evidence of differential expression by sex.

In addition, we also filtered our dataset by 155 effector genes linked to CRC risk, which have been recently published (Fernandez-Rozadilla *et al*., 2022). Genes *BMP5* and *LRIG1* showed a significant adjusted p-value below 0.05, however the log2 fold-change was small in both cases (−0.22 and 0.14). The remaining genes did not reach significance (**Supplementary Table 3**).

A single nucleotide polymorphism (SNP) on gene *ERBB4* was previously described as the most significant SNP associated with overall survival (HR = 1.24, 95%CI = 1.16-1.32, P = 1.9 × 10-7) (Wills *et al*., 2021). We therefore further wanted to understand whether this gene showed some effect in our dataset by sex. However, no significance of this gene was observed in our results.

### Association between BMI and Gene Expression

DEA for BMI was performed by combining both sexes in one model and using sex as a covariate. Stratified models for each sex have been built as well (**Table 3**).

**TABLE 3.**
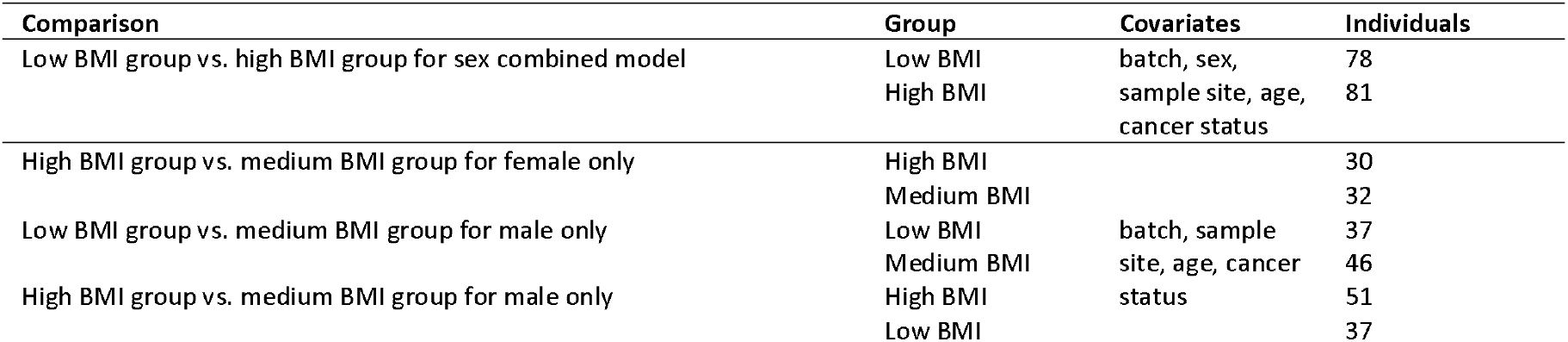
Overview Analysis of Association between BMI and Gene Expression. Table only shows models with significant results.

The combined model showed one significant gene, *TXLNGY*, for the comparison between samples with low and high BMI **(Figure 2)**. DEA for females only, identified the gene *MTND2P28* for BMI group high versus medium **(Figure 2**). The models comparing the BMI groups for males showed gene *CPB1* in the low versus medium comparison and genes *IGKV2-29* and *H19* for the comparison high versus medium (**Figure 2)**. Overall, we found a limited number of significant genes. A detailed table can be found in the Supplementary Material (**Supplementary Table 2**).

**FIGURE 2.**
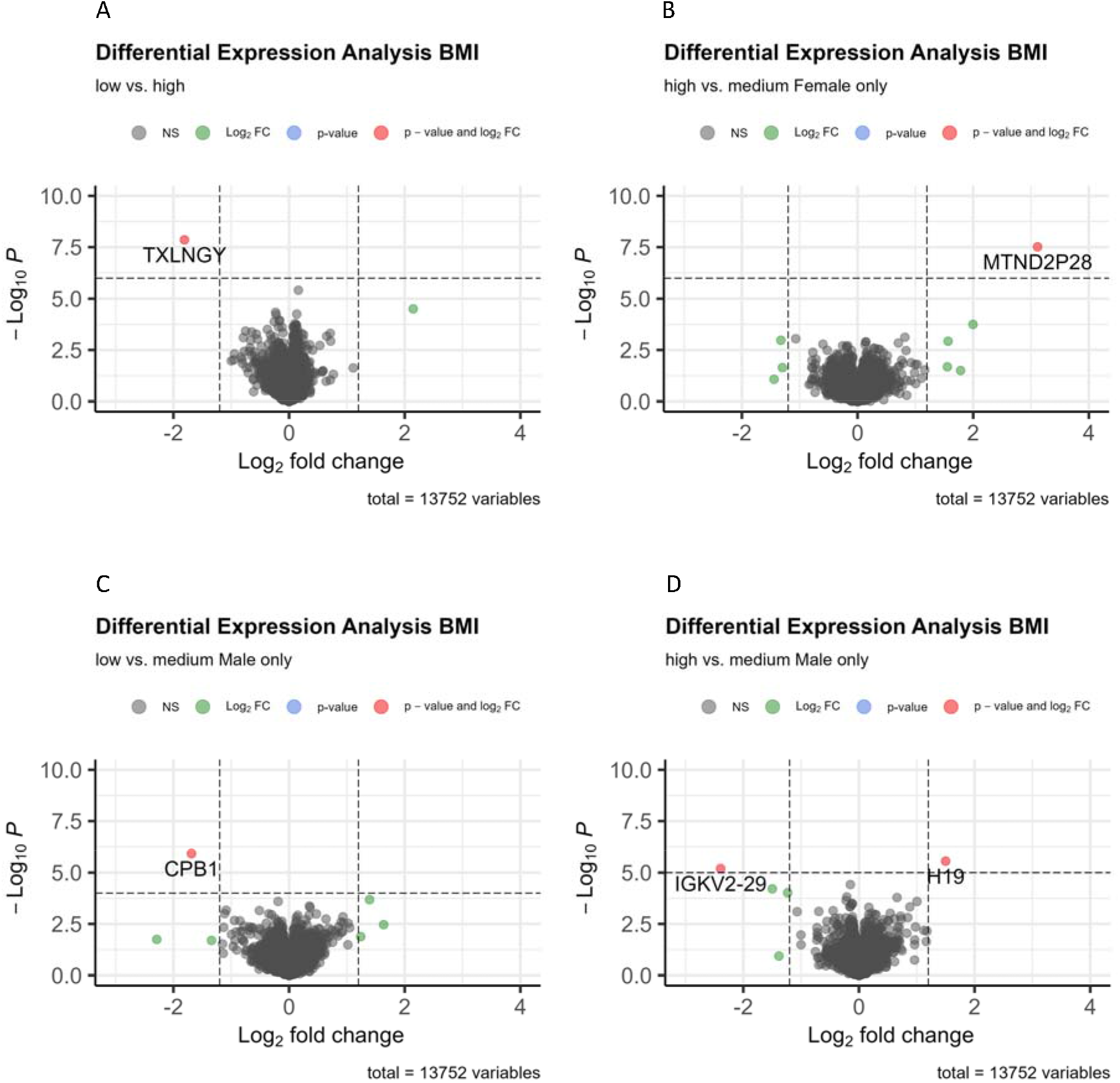
DEA by BMI. Differentially downregulated and upregulated genes expression in **A** low vs. high BMI, in **B** high vs. medium BMI (female only), in **C** low vs. medium BMI (male only) and in **D** high vs. medium BMI (male only) were identified using volcano plots.

We wanted to further explore the hypothesis that CRC risk genes might be affected by BMI. We repeated the analysis by specifically analysing 46 candidate target genes, which have been identified previously (Law *et al*., 2019). None of the genes reached nominal significance. Finally, we wanted to understand whether genes on the sex chromosomes could be influenced by BMI. We enhanced the analysis by only focusing on Y (18 genes) and/or X (462 genes) chromosome genes for BMI. Again, no significant genes were identified.

Finally, we also limited our dataset to 155 effector genes linked to CRC risk (Fernandez-Rozadilla *et al*., 2022), however none of the remaining 126 genes in our dataset showed a significant adjusted p-value.

### Association between Cancer Status and Gene Expression

In order to understand which role an ongoing cancer condition could have on the gene expression of normal colorectal epithelium, normal colorectal mucosa of individuals without cancer was compared to normal colorectal tissue of individuals with cancer, neoplasia and adenomas (**Table 4**). 100 out of our 365 patients had cancer at the time of normal colorectal tissue collection, while 68 of 365 individuals had an adenoma.

**TABLE 4.** Overview Analysis of Association between Cancer Status and Gene Expression

| Comparison | Group | Covariates | Individuals in SOCCS | Individuals in INTERMPHEN |
| --- | --- | --- | --- | --- |
| No cancer vs. with cancer | Without cancer |  | 109 |  |
|  | With cancer |  | 100 |  |
| No cancer vs. with neoplasia | Without cancer | batch, sample site, sex, BMI, age | 109 |  |
|  | With neoplasia |  | 110 |  |
| No cancer vs. with adenoma | Without cancer |  | 109 | 69 |
|  | With adenoma |  | 10 | 58 |

Three differentially expressed genes (*H3C15, FOSB* and *TFF2*) have been identified when comparing individuals with no cancer with individuals with cancer (**Table 5**). The genes *H3C15, FOSB* and *TFF2* were significant when comparing individuals without cancer with individuals with neoplasia in SOCCS (**Table 5**). Finally, gene *ZFAT* was significant when comparing individuals without cancer with individuals with adenomas (**Table 5**). Since INTERMPHEN does not contain cancer samples, we repeated only the last comparison, which showed no significance (**Table 5**).

**TABLE 5.**
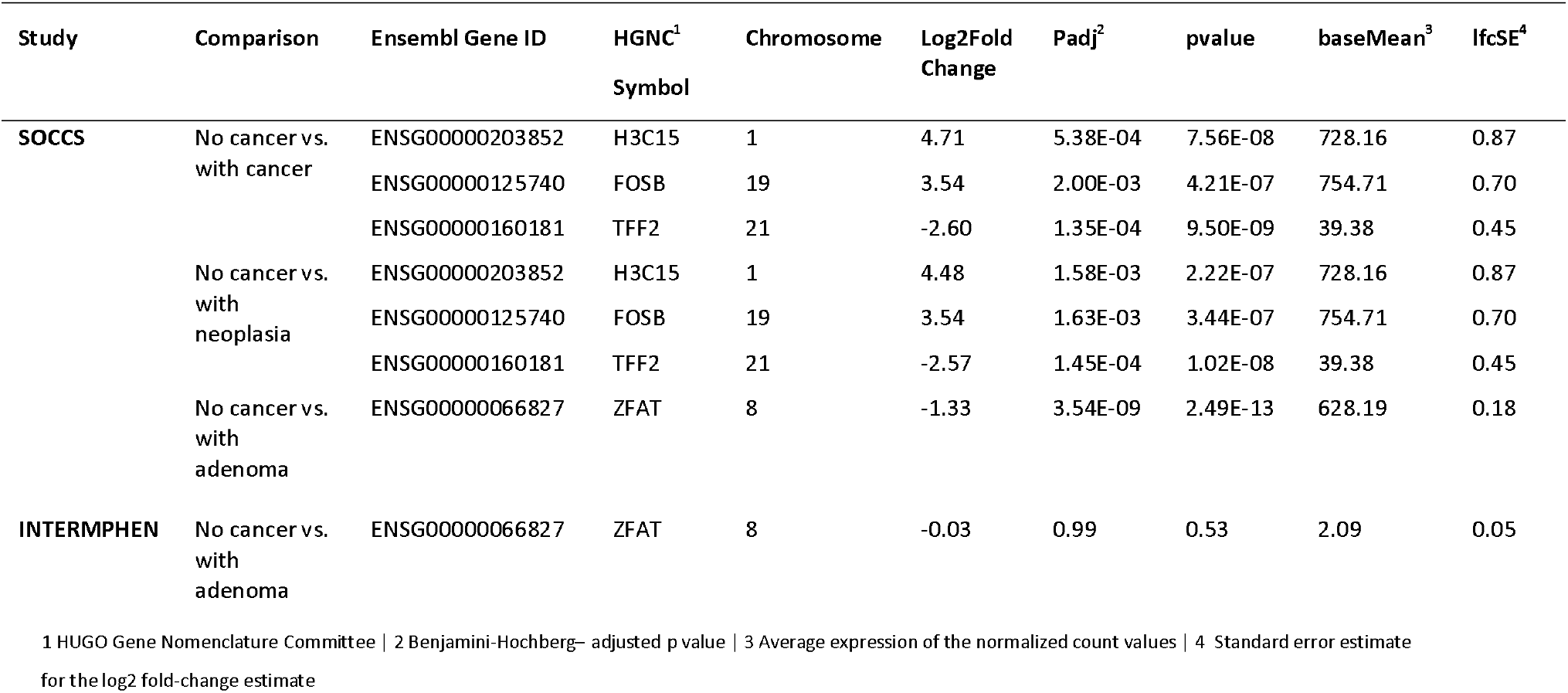
Results of DEA of **cancer status** comparisons in SOCCS and INTERMPHEN. Models have been adjusted for batch, sample site, sex, BMI and age. Included genes with adjusted p-value below 0.05 and log2 fold-change threshold above 1.2.

## DISCUSSION

In this exploratory analysis of normal colorectal epithelium, we performed RNA sequencing of 365 individuals to examine the effect of sex, BMI and cancer status on gene expression.

DEA identified two genes which were not located on the X or Y chromosome and fulfilled our filtering requirements. The volcano plot, which excluded all sex chromosomes, shows that more genes with p-value below 0.05 can be identified, however they were excluded because of the chosen log2 fold-change threshold (**Supplementary Figure 1**). Since none of the effects could be replicated in a second dataset, our findings suggest that there is no strong association between sex and gene expression in normal colorectal mucosa tissue. The limited reliability of gene *IGHV2-70D*, due to an immunoglobulin heavy chain, needs to be also taken into consideration.

We further explored the hypothesis that expression of CRC risk genes might be affected by sex by limiting our dataset to 46 candidate target genes (Law *et al*., 2019). No significant genes were found (**Supplementary Table 1**). The highest log2 fold-change was 0.24 and the smallest adjusted p-value was 0.32.

Furthermore, the 155 effector genes linked to CRC risk, as described previously, did also not reach significance in our dataset (**Supplementary Table 3**). Finally, the gene *ERBB4*, which SNP was reported as the most significant SNP associated with overall survival, did also not show significance in our results.

Recent publications have analysed the impact of sex on gene expression in transverse colon (samples from 136 females and 232 males) as well as sigmoid colon (samples from 113 female and 205 male) (Oliva et al., 2020). In total, 3262 sex-differentially expressed genes were reported for transverse colon as well as 3510 for sigmoid colon. We were able to identify 11 genes not located on the sex chromosomes (*FRG1CP, KRBOX1, NLRP2, FAM24B, MTLN, FRG1BP, ZMYND11, DUOX2, SLFN13, GPR15* and *MIR4458HG)* in our discovery dataset results, which overlapped with the top 500 Oliva et al. findings for sigmoid colon (**Supplementary Table 4**). Although all of our genes presented an adjusted p-value below 0.05, the log2 fold-change showed an effect below our cut-off of 1.2. We conducted a Hypergeometric Distribution Test (using the function phyper in R) comparing the proportions of our identified significant genes in the SOCCS gene pool compared to the proportion in the top 500 differently expressed genes in the Oliva et al. publication (R: The Hypergeometric Distribution, 2022).

We concluded that the observed proportions in the top 500 genes are significantly different from the expected proportions (p-value =3.60e-16) and we therefore report a significant enrichment in our dataset (**Supplementary Figure 2**). We repeated the analysis for the top 500 expressed genes in transverse colon and compared the overlap with our discovery dataset. We identified 9 genes, which overlapped with the previous genes of sigmoid colon, and we found two additional genes (*MOAP1* and *MYEF2*) (**Supplementary Table 5**). The Hypergeometric Distribution Test showed again a significant enrichment in our dataset. Finally, we wanted to understand whether our p-value decreases even more if we minimise our dataset by setting the p-value threshold to 0.01 and 0.001. Interestingly, the more we restrict our dataset and then compare it to the external dataset, the more our p-value of the Hypergeometric Distribution Test decreases (with p-value=0.01 we receive a p-value of 5.89e-26 and with p-value=0.001 it is 1.73936e-33). We also compared our results with 51 transverse colon and 45 sigmoid colon sex-biased differentially expressed genes (Lopes-Ramos et al., 2020) and we could not see common genes with our results beside some matches in the sex chromosomes. It should be highlighted that the majority of current studies in this field use the GTEx dataset (Lopes-Ramos et al., 2020) (Oliva et al., 2020) (Naqvi et al., 2019) (Melé et al., 2015) (Gershoni and Pietrokovski, 2017). If another normal colorectal mucosa dataset than GTEx is used, it tends to be very small in sample size in comparison to the datasets presented in our study (Hases et al., 2021). Moreover, the presented dataset of our paper is completely independent from GTEx.

The analysis of BMI was performed in two steps: While in step one males and females were combined in one model, in step two males and females were examined separately. It should be noted that this analysis was only run on one dataset (SOCCS), since the second dataset did not include BMI. The models for males and females separately identified slightly more genes than a combined model of both sexes. The volcano plot shows that apart from the five identified genes no genes have a significant p-value (**Figure 2**), although some would pass the log2 fold-change cut-off.

Recent efforts compared gene expression between 90 normal and overweight individuals and identified seven differentially expressed genes (*RBM20, SEP12, AX748233, SLC30A3, WTIP, CASP10* and *OR12D3*) associated with BMI (Joseph *et al*., 2019). Only the gene *CASP10 overlapped with our* results but with a log2 fold-change of 0.04 and an adjusted p-value of 1. In addition, we further examined the hypothesis that the expression of CRC risk genes might be affected by BMI by limiting our dataset to 46 candidate target genes (Law *et al*., 2019). No significant genes were identified. The analysis was further enhanced by limiting the dataset to the genes located on the X and Y chromosome only, which again did not show significant genes. Finally, the hypothesis that the gene expression of 155 effector genes linked to CRC risk might be influenced by BMI was rejected.

Besides our interest in the association between sex and BMI and gene expression we wanted to understand which role an ongoing cancer condition could have on the gene expression of normal colorectal epithelium. As previously mentioned, most studies in the past focused on comparing tumour tissue and adjacent normal tissue. The comparison of only normal tissue between patients with/without cancer is rarely performed and we aimed to ensure that gene expression of a normal colorectal tissue is not affected by a cancer condition. Only a few genes (*H3C15, FOSB, TFF2* and *ZFAT*) were identified in the DEA of samples of individuals without cancer against samples of individuals with an ongoing cancer condition/neoplasia or adenoma. While *H3C15, FOSB* and *ZFAT* have not been classified as cancer-related genes, *TFF2* have been reported to be related to cancer (Search: *Tff2* - *The Human Protein Atlas*, 2022, p. 2). A high expression of *TFF2* has been associated with unfavourable outcome in liver cancer. *FOSB, ZFAT* and *TFF2* show differentially expression in cancer while *H3C15* does not (Search: *Tff2* - *The Human Protein Atlas*, 2022, p. 2).

Finally, our method of analysis has some limitations. Even though we tried to eliminate any confounding factors in our analysis by adjusting the models accordingly, we cannot completely rule out any effects of unequal distributions of individuals with/without ongoing cancer condition as well as effects due to cases taken from different batches. In order to ensure similar grouping of individuals in BMI groups we decided to use tertiles for the BMI groups instead of following the traditional grouping of underweight/normal/overweight/obese 1/obese 2. It would be interesting to replicate the analysis with these grouping in a larger dataset, which would have sufficient cases for each subgroup. Moreover, we need to highlight that our BMI comparisons involved many comparisons between different groups, which might influence our results. A further limitations of our study might be the chosen log2 fold threshold of 1.2. Other studies have chosen lower cut-offs and therefore presented more differentially expressed genes. Apart from this, it should be noted that we used two different methods for the two datasets due to the limitations of DESEQ2 to analyse repeating measurements. We followed the established workflows for the different algorithms, which resulted in e.g. different ways of normalising the data. The findings between both datasets are therefore not directly comparable and should be interpreted with caution. However, it remains unclear why our DEA in SOCCS identifies effects which cannot be found at all in INTERMPHEN. This rather contradictory result may be simply due to the fact that no relationship can be found between sex, BMI or cancer status and gene expression.

## CONCLUSION

We tried to answer the question whether sex and BMI are associated with gene expression in normal colorectal tissue. Our analysis was enhanced by evaluating possible confounding effects of cancer presence on the gene expression. Only a limited number of genes for sex, BMI and confounding by cancer status were identified via DEA, which showed some trend towards differential expression. However, the findings could not be replicated in a second dataset and did not withstand multiple testing.

We therefore cannot observe strong systematic differences between sex, BMI group and cancer status on gene expression in normal colorectal epithelium from human subjects. However, sample size and other confounding factors might limit our analysis. Also it needs to be highlighted that we observed a significant enrichment of previously identified genes in our dataset, which however did not withstand our filtering requirements. In spite of its limitations, the findings of this study suggest that an increased risk by male and obese individuals cannot be explained in gene expression differences in sex and BMI. Future studies to validate our findings are recommended.

## Acknowledgments

The work was supported by Cancer Research UK program grants to MGD and SMF (DRCPGM\100012 and C348/A18927). There was also support for funding of the infrastructure and staffing of the Edinburgh CRUK Cancer Research Centre. MGD also received Medical Research Council grant funding as MRC Programme Leader within the MRC Human Genetics Unit (U127527198).

Prof. Evropi Theodoratou is supported by a Cancer Research UK Career Development Fellowship (C31250/A22804).

## Notes

### Competing Interest Statement

The authors have declared no competing interest.

### Summary of Updates

Adding acknowledgments.

## References

‘Analysis pipelines for the GTEx Consortium and TOPMed’ (2022). Broad Institute. Available at: https://github.com/broadinstitute/gtex-pipeline/blob/8c0b763cfe35d5a15c88ca8ca91c73fa0a5cfc52/rnaseq/README.md (Accessed: 1 August 2020).

Bardou, M., Barkun, A.N. and Martel, M. (2013) ‘Obesity and colorectal cancer’, Gut, 62((6), pp. 933–947. Available at: https://doi.org/10.1136/gutjnl-2013-304701.

Benjamini, Y. and Hochberg, Y. (1995) ‘Controlling the False Discovery Rate: A Practical and Powerful Approach to Multiple Testing’, Journal of the Royal Statistical Society. Series B (Methodological), 57((1), pp. 289–300.

Bray, F. et al. (2018) ‘Global cancer statistics 2018: GLOBOCAN estimates of incidence and mortality worldwide for 36 cancers in 185 countries’, CA: a cancer journal for clinicians, 68((6), pp. 394–424. Available at: https://doi.org/10.3322/caac.21492.

Dai, Z., Xu, Y.-C. and Niu, L. (2007) ‘Obesity and colorectal cancer risk: a meta-analysis of cohort studies’, World Journal of Gastroenterology, 13((31), pp. 4199–4206. Available at: https://doi.org/10.3748/wjg.v13.i31.4199.

DeLuca, D.S. et al. (2012) ‘RNA-SeQC: RNA-seq metrics for quality control and process optimization’, Bioinformatics (Oxford, England), 28((11), pp. 1530–1532. Available at: https://doi.org/10.1093/bioinformatics/bts196.

Dobin, A. et al. (2013) ‘STAR: ultrafast universal RNA-seq aligner’, Bioinformatics (Oxford, England), 29((1), pp. 15–21. Available at: https://doi.org/10.1093/bioinformatics/bts635.

dupcor: Correlation Between Duplicates in limma: Linear Models for Microarray Data (2022). Available at: https://rdrr.io/bioc/limma/man/dupcor.html (Accessed: 25 September 2022).

Fernandez-Rozadilla, C. et al. (2018) ‘Telomere length and genetics are independent colorectal tumour risk factors in an evaluation of biomarkers in normal bowel’, British Journal of Cancer, 118((5), pp. 727–732. Available at: https://doi.org/10.1038/bjc.2017.486.

Fernandez-Rozadilla, C. et al. (2022) ‘Deciphering colorectal cancer genetics through multi-omic analysis of 100,204 cases and 154,587 controls of European and East Asian ancestries’. Zenodo. Available at: https://doi.org/10.5281/zenodo.6472285.

GENCODE - Human Release 26 (2020). Available at: https://www.gencodegenes.org/human/release_26.html (Accessed: 1 August 2020).

Gershoni, M. and Pietrokovski, S. (2017) ‘The landscape of sex-differential transcriptome and its consequent selection in human adults’, BMC biology, 15((1), p. 7. Available at: https://doi.org/10.1186/s12915-017-0352-z.

Gordon K. Smyth et al. (2021) Linear Models for Microarray and RNA-Seq Data. Available at: https://www.bioconductor.org/packages/devel/bioc/vignettes/limma/inst/doc/usersguide.pdf(Accessed: 10 September 2022).

GTEx Portal (2022). Available at: https://gtexportal.org/home/ (Accessed: 1 September 2022).

Hases, L. et al. (2021) ‘The Importance of Sex in the Discovery of Colorectal Cancer Prognostic Biomarkers’, International Journal of Molecular Sciences, 22((3), p. 1354. Available at: https://doi.org/10.3390/ijms22031354.

Huang, X., Stern, D.F. and Zhao, H. (2016) ‘Transcriptional Profiles from Paired Normal Samples Offer Complementary Information on Cancer Patient Survival – Evidence from TCGA Pan-Cancer Data’, Scientific Reports, 6((1), p. 20567. Available at: https://doi.org/10.1038/srep20567.

Joseph, P. et al. (2019) ‘Comprehensive and Systematic Analysis of Gene Expression Patterns Associated with Body Mass Index’, Scientific Reports, 9. Available at: https://doi.org/10.1038/s41598-019-43881-5.

Kim, H. and Giovannucci, E.L. (2017) ‘Sex differences in the association of obesity and colorectal cancer risk’, Cancer causes & control: CCC, 28((1), pp. 1–4. Available at: https://doi.org/10.1007/s10552-016-0831-5.

Law, P.J. et al. (2019) ‘Association analyses identify 31 new risk loci for colorectal cancer susceptibility’, Nature Communications, 10((1), p. 2154. Available at: https://doi.org/10.1038/s41467-019-09775-w.

Li, B. and Dewey, C.N. (2011) ‘RSEM: accurate transcript quantification from RNA-Seq data with or without a reference genome’, BMC Bioinformatics, 12((1), p. 323. Available at: https://doi.org/10.1186/1471-2105-12-323.

Loo, L.W.M., Lemire, M. and Le Marchand, L. (2017) ‘In silico pathway analysis and tissue specific cis-eQTL for colorectal cancer GWAS risk variants’, BMC genomics, 18((1), p. 381. Available at: https://doi.org/10.1186/s12864-017-3750-2.

Lopes-Ramos, C.M. et al. (2020) ‘Sex Differences in Gene Expression and Regulatory Networks across 29 Human Tissues’, Cell Reports, 31((12), p. 107795. Available at: https://doi.org/10.1016/j.celrep.2020.107795.

Love, M. et al. (2022) ‘DESeq2: Differential gene expression analysis based on the negative binomial distribution’. Bioconductor version: Release (3.15). Available at: https://doi.org/10.18129/B9.bioc.DESeq2.

Love, M.I., Huber, W. and Anders, S. (2014) ‘Moderated estimation of fold change and dispersion for RNA-seq data with DESeq2’, Genome Biology, 15((12), p. 550. Available at: https://doi.org/10.1186/s13059-014-0550-8.

Melé, M. et al. (2015) ‘Human genomics. The human transcriptome across tissues and individuals’, Science (New York, N.Y.), 348((6235), pp. 660–665. Available at: https://doi.org/10.1126/science.aaa0355.

Michael I. Love, S.A., Vladislav Kim and Wolfgang Huber (2019) RNA-seq workflow: gene-level exploratory analysis and differential expression. Available at: https://bioconductor.org/packages/release/workflows/vignettes/rnaseqGene/inst/doc/rnaseqGene.html (Accessed: 6 July 2021).

Naqvi, S. et al. (2019) ‘Conservation, acquisition, and functional impact of sex-biased gene expression in mammals’, Science (New York, N.Y.), 365((6450), p. eaaw7317. Available at: https://doi.org/10.1126/science.aaw7317.

Oliva, M. et al. (2020) ‘The impact of sex on gene expression across human tissues’, Science, 369((6509), p. eaba3066. Available at: https://doi.org/10.1126/science.aba3066.

R: The Hypergeometric Distribution (2022). Available at: https://stat.ethz.ch/R-manual/R-devel/library/stats/html/Hypergeometric.html (Accessed: 25 October 2022).

R: The R Project for Statistical Computing (2022). Available at: https://www.r-project.org/ (Accessed: 19 October 2022).

Ritchie, M.E. et al. (2015) ‘limma powers differential expression analyses for RNA-sequencing and microarray studies’, Nucleic Acids Research, 43((7), p. e47. Available at: https://doi.org/10.1093/nar/gkv007.

Search: Tff2 - The Human Protein Atlas (2022). Available at: https://www.proteinatlas.org/search/Tff2 (Accessed: 19 October 2022).

Tarasiuk, A., Mosińska, P. and Fichna, J. (2018) ‘The mechanisms linking obesity to colon cancer: An overview’, Obesity Research & Clinical Practice, 12((3), pp. 251–259. Available at: https://doi.org/10.1016/j.orcp.2018.01.005.

Vaughan-Shaw, Peter.G. et al. (2021) ‘Differential genetic influences over colorectal cancer risk and gene expression in large bowel mucosa’, International Journal of Cancer, 149((5), pp. 1100–1108. Available at: https://doi.org/10.1002/ijc.33616.

Vaughan-Shaw, P.G. et al. (2022) ‘Factors influencing patterns of gene expression in large bowel mucosa’. bioRxiv, p. 2022.08.30.505238. Available at: https://doi.org/10.1101/2022.08.30.505238.

Wills, C. et al. (2021) ‘A genome-wide search for determinants of survival in 1926 patients with advanced colorectal cancer with follow-up in over 22,000 patients’, European Journal of Cancer, 159, pp. 247–258. Available at: https://doi.org/10.1016/j.ejca.2021.09.047.

World Cancer Report: Cancer Research for Cancer Prevention (PDF) (2022) IARC E-Bookshop. Available at: https://shop.iarc.fr/products/world-cancer-report-cancer-research-for-cancer-prevention-pdf (Accessed: 25 September 2022).

