## Supplementary Material for "Exploring the association between Body Mass Index, Sex and Gene Expression in human colorectal epithelium"

Principal component analysis (PCA) was used to identify outliers and to check whether samples cluster together. Three outliers have been excluded for SOCCS, leaving 238 samples for further analysis. Clustering of samples by batch, sample site and sex was identified in the PCA Analysis for SOCCS and INTERMPHEN therefore the models were controlled for these factors where relevant.

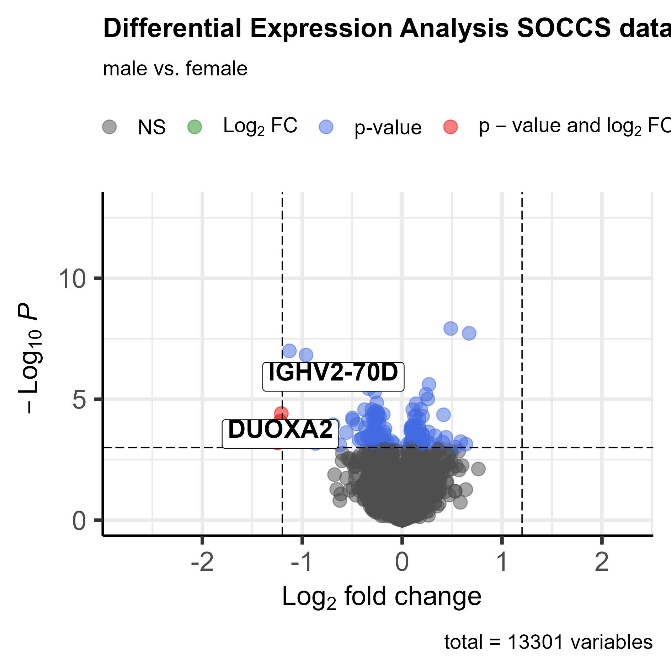

**FIGURE 1** | Distribution of genes of Differential Expression Analysis (DEA) male vs. female. Included samples from SOCCS dataset**.**

**TABLE 1** | Results of DEA **male vs. female in 46 candidate target genes**. Models have been adjusted for batch, sample site, BMI, age and cancer status. Included samples from SOCCS Study.

^1 HUGO Gene Nomenclature Committee | 2 Benjamini-Hochberg– adjusted p value | 3 Average expression of the normalized count values | 4^ ^Standard error estimate for the log2 fold change estimate^

| **Ensembl Gene ID** | **HGNC^1^ Symbol** | **Chromosome** | | **Log2Fold Change** | | **Padj^2^** | | **Pvalue** | | **baseMean^3^** | | **lfcSE^4^** | |
| --- | --- | --- | --- | --- | --- | --- | --- | --- | --- | --- | --- | --- | --- |
| ENSG00000033800 | PIAS1 | | 15 | | -0.02 | | 0.77 | | 0.44 | | 2228.98 | | 0.03 |
| ENSG00000050405 | LIMA1 | | 12 | | -0.02 | | 0.83 | | 0.63 | | 15446.95 | | 0.04 |
| ENSG00000057657 | PRDM1 | | 6 | | -0.03 | | 0.83 | | 0.68 | | 1122.01 | | 0.07 |
| ENSG00000066084 | DIP2B | | 12 | | -0.02 | | 0.63 | | 0.30 | | 5259.73 | | 0.02 |
| ENSG00000066117 | SMARCD1 | | 12 | | -0.01 | | 0.83 | | 0.65 | | 977.99 | | 0.02 |
| ENSG00000096060 | FKBP5 | | 6 | | -0.03 | | 0.83 | | 0.67 | | 2901.04 | | 0.08 |
| ENSG00000110777 | POU2AF1 | | 11 | | -0.18 | | 0.36 | | 0.05 | | 413.42 | | 0.09 |
| ENSG00000111252 | SH2B3 | | 12 | | -0.07 | | 0.46 | | 0.14 | | 1136.76 | | 0.04 |
| ENSG00000115085 | ZAP70 | | 2 | | -0.21 | | 0.32 | | 0.02 | | 197.55 | | 0.09 |
| ENSG00000116209 | TMEM59 | | 1 | | 0.05 | | 0.62 | | 0.25 | | 15100.70 | | 0.04 |
| ENSG00000118971 | CCND2 | | 12 | | 0.06 | | 0.52 | | 0.17 | | 11638.59 | | 0.05 |
| ENSG00000120693 | SMAD9 | | 13 | | 0.14 | | 0.32 | | 0.02 | | 800.72 | | 0.06 |
| ENSG00000123384 | LRP1 | | 12 | | -0.01 | | 0.91 | | 0.85 | | 11873.50 | | 0.05 |
| ENSG00000124198 | ARFGEF2 | | 20 | | 0.03 | | 0.46 | | 0.11 | | 8933.48 | | 0.02 |
| ENSG00000125378 | BMP4 | | 14 | | -0.14 | | 0.36 | | 0.04 | | 466.91 | | 0.07 |
| ENSG00000125845 | BMP2 | | 20 | | 0.00 | | 0.96 | | 0.93 | | 1549.25 | | 0.05 |
| ENSG00000130702 | LAMA5 | | 20 | | 0.10 | | 0.62 | | 0.26 | | 699.93 | | 0.09 |
| ENSG00000135111 | TBX3 | | 12 | | -0.03 | | 0.83 | | 0.61 | | 400.66 | | 0.06 |
| ENSG00000135862 | LAMC1 | | 1 | | 0.01 | | 0.91 | | 0.84 | | 4431.44 | | 0.06 |
| ENSG00000135926 | TMBIM1 | | 2 | | -0.04 | | 0.74 | | 0.39 | | 8134.51 | | 0.04 |
| ENSG00000136122 | BORA | | 13 | | 0.04 | | 0.74 | | 0.39 | | 480.24 | | 0.05 |
| ENSG00000136205 | TNS3 | | 7 | | -0.04 | | 0.62 | | 0.27 | | 5318.91 | | 0.03 |
| ENSG00000136997 | MYC | | 8 | | 0.12 | | 0.46 | | 0.08 | | 1186.36 | | 0.07 |
| ENSG00000143507 | DUSP10 | | 1 | | 0.00 | | 0.97 | | 0.97 | | 202.92 | | 0.06 |
| ENSG00000147883 | CDKN2B | | 9 | | -0.08 | | 0.62 | | 0.26 | | 3384.63 | | 0.08 |
| ENSG00000148737 | TCF7L2 | | 10 | | -0.07 | | 0.46 | | 0.11 | | 1722.12 | | 0.04 |
| ENSG00000149679 | CABLES2 | | 20 | | 0.01 | | 0.91 | | 0.86 | | 200.58 | | 0.04 |
| ENSG00000162552 | WNT4 | | 1 | | -0.11 | | 0.46 | | 0.12 | | 162.67 | | 0.07 |
| ENSG00000163935 | SFMBT1 | | 3 | | 0.02 | | 0.83 | | 0.57 | | 1413.59 | | 0.04 |
| ENSG00000164161 | HHIP | | 4 | | 0.03 | | 0.83 | | 0.66 | | 1239.76 | | 0.06 |
| ENSG00000166923 | GREM1 | | 15 | | 0.24 | | 0.37 | | 0.06 | | 1893.75 | | 0.12 |
| ENSG00000168036 | CTNNB1 | | 3 | | -0.01 | | 0.83 | | 0.68 | | 18409.04 | | 0.03 |
| ENSG00000168769 | TET2 | | 4 | | 0.01 | | 0.83 | | 0.67 | | 4099.60 | | 0.02 |
| ENSG00000176920 | FUT2 | | 19 | | -0.09 | | 0.46 | | 0.13 | | 1778.08 | | 0.06 |
| ENSG00000178449 | COX14 | | 12 | | 0.11 | | 0.32 | | 0.01 | | 622.96 | | 0.04 |
| ENSG00000183431 | SF3A3 | | 1 | | 0.03 | | 0.63 | | 0.28 | | 1864.66 | | 0.03 |
| ENSG00000196167 | COLCA1 | | 11 | | -0.07 | | 0.78 | | 0.47 | | 463.13 | | 0.10 |
| ENSG00000196396 | PTPN1 | | 20 | | -0.01 | | 0.84 | | 0.72 | | 1651.00 | | 0.03 |
| ENSG00000214290 | COLCA2 | | 11 | | 0.02 | | 0.91 | | 0.82 | | 149.27 | | 0.10 |
| ENSG00000240498 | CDKN2B-AS1 | | 9 | | 0.08 | | 0.77 | | 0.43 | | 923.59 | | 0.10 |

**TABLE 2 |** Results of DEA for **BMI group** comparisons. Models have been adjusted for batch, sex, sample site, age and cancer status. Included samples from SOCCS. Included genes with adjusted p-value below 0.05 and log2fold change threshold above 1.2.

| **Comparison** | **Ensembl Gene ID** | **HGNC^1^**  **Symbol** | **Chromosome** | **Log2Fold Change** | **Padj^2^** | **pvalue** | **baseMean^3^** | **lfcSE^4^** |
| --- | --- | --- | --- | --- | --- | --- | --- | --- |
| low vs. high | ENSG00000131002 | TXLNGY | Y | -1.81 | 6.75E-05 | 1.39E-08 | 2825.65 | 0.32 |
| high vs. medium female only | ENSG00000225630 | MTND2P28 | 1 | 3.12 | 4.32E-04 | 2.97E-08 | 498.23 | 0.56 |
| low vs. medium male only | ENSG00000153002 | CPB1 | 3 | -1.69 | 0.02 | 1.15E-06 | 257.03 | 0.35 |
| high vs. medium male only | ENSG00000253998 | IGKV2-29 | 2 | -2.40 | 0.04 | 6.14E-06 | 119.86 | 0.53 |
|  | ENSG00000130600 | H19 | 11 | 1.49 | 0.04 | 4.04E-06 | 82.56 | 0.32 |

^1 HUGO Gene Nomenclature Committee | 2 Benjamini-Hochberg– adjusted p value | 3 Average expression of the normalized count values | 4^ ^Standard error estimate for the log2 fold change estimate^

**TABLE 3** | Results of DEA **male vs. female in top 30 of 155 effector genes by adjusted p-value**. Models have been adjusted for batch, sample site, BMI, age and cancer status. Included samples from SOCCS Study.

^1 HUGO Gene Nomenclature Committee | 2 Benjamini-Hochberg– adjusted p value | 3 Average expression of the normalized count values | 4^ ^Standard error estimate for the log2 fold change estimate^

| **Ensembl Gene ID** | **HGNC^1^ Symbol** | **Chromosome** | **Log2Fold Change** | **Padj^2^** | **Pvalue** | **baseMean^3^** | **lfcSE^4^** |
| --- | --- | --- | --- | --- | --- | --- | --- |
| ENSG00000112175 | BMP5 | 6 | -0.22 | 0.01 | 5.21E-05 | 621.03 | 0.06 |
| ENSG00000144749 | LRIG1 | 3 | 0.14 | 0.01 | 9.14E-05 | 2712.22 | 0.04 |
| ENSG00000149591 | TAGLN | 11 | 0.47 | 0.08 | 1.94E-03 | 5007.70 | 0.15 |
| ENSG00000124126 | PREX1 | 20 | -0.16 | 0.10 | 3.17E-03 | 1115.32 | 0.05 |
| ENSG00000139636 | LMBR1L | 12 | 0.08 | 0.13 | 5.28E-03 | 753.28 | 0.03 |
| ENSG00000163431 | LMOD1 | 1 | 0.38 | 0.14 | 6.55E-03 | 1237.31 | 0.14 |
| ENSG00000183579 | ZNRF3 | 22 | 0.12 | 0.15 | 8.23E-03 | 543.98 | 0.05 |
| ENSG00000178573 | MAF | 16 | -0.15 | 0.15 | 9.45E-03 | 1017.98 | 0.06 |
| ENSG00000095739 | BAMBI | 10 | -0.25 | 0.20 | 1.88E-02 | 214.66 | 0.11 |
| ENSG00000125378 | BMP4 | 14 | -0.16 | 0.20 | 1.80E-02 | 468.94 | 0.07 |
| ENSG00000125827 | TMX4 | 20 | 0.09 | 0.20 | 2.03E-02 | 1777.68 | 0.04 |
| ENSG00000144857 | BOC | 3 | 0.22 | 0.20 | 1.51E-02 | 228.10 | 0.09 |
| ENSG00000178449 | COX14 | 12 | 0.10 | 0.20 | 1.66E-02 | 620.38 | 0.04 |
| ENSG00000088766 | CRLS1 | 20 | 0.06 | 0.22 | 2.63E-02 | 1732.75 | 0.03 |
| ENSG00000198876 | DCAF12 | 9 | 0.04 | 0.22 | 2.60E-02 | 1777.68 | 0.02 |
| ENSG00000064270 | ATP2C2 | 16 | 0.09 | 0.28 | 3.57E-02 | 5783.73 | 0.04 |
| ENSG00000120693 | SMAD9 | 13 | 0.12 | 0.28 | 3.73E-02 | 796.16 | 0.06 |
| ENSG00000113638 | TTC33 | 5 | 0.07 | 0.32 | 4.52E-02 | 739.76 | 0.03 |
| ENSG00000079819 | EPB41L2 | 6 | -0.09 | 0.34 | 7.52E-02 | 4474.42 | 0.05 |
| ENSG00000113719 | ERGIC1 | 5 | 0.05 | 0.34 | 6.59E-02 | 7581.42 | 0.03 |
| ENSG00000133121 | STARD13 | 13 | 0.07 | 0.34 | 7.01E-02 | 1294.70 | 0.04 |
| ENSG00000136270 | TBRG4 | 7 | 0.06 | 0.34 | 6.25E-02 | 1900.67 | 0.03 |
| ENSG00000142632 | ARHGEF19 | 1 | -0.10 | 0.34 | 6.80E-02 | 146.06 | 0.06 |
| ENSG00000146833 | TRIM4 | 7 | 0.04 | 0.34 | 6.51E-02 | 1142.02 | 0.02 |
| ENSG00000148737 | TCF7L2 | 10 | -0.08 | 0.34 | 6.37E-02 | 1733.15 | 0.04 |
| ENSG00000163933 | RFT1 | 3 | 0.04 | 0.34 | 7.59E-02 | 709.31 | 0.02 |
| ENSG00000166923 | GREM1 | 15 | 0.22 | 0.34 | 7.71E-02 | 1887.37 | 0.12 |
| ENSG00000167693 | NXN | 17 | 0.19 | 0.34 | 6.37E-02 | 172.02 | 0.10 |
| ENSG00000234608 | MAPKAPK5-AS1 | 12 | 0.07 | 0.34 | 5.43E-02 | 650.85 | 0.04 |
| ENSG00000111252 | SH2B3 | 12 | -0.08 | 0.35 | 9.18E-02 | 1143.56 | 0.05 |

**TABLE 4|** Differentially expressed genes with adjusted p-value below 0.05 overlapping with top 500 differentially expressed genes in sigmoid colon in external gene list (Oliva *et al.*, 2020)**.**

| **Ensembl Gene ID** | **HGNC^1^**  **Symbol** | **Chromosome** | **Log2Fold Change** | **Padj^2^** | **pvalue** |
| --- | --- | --- | --- | --- | --- |
| ENSG00000282826 | FRG1CP | 20 | 0.49 | 3.26E-06 | 1.12E-08 |
| ENSG00000240747 | KRBOX1 | 3 | 0.67 | 5.50E-06 | 1.97E-08 |
| ENSG00000022556 | NLRP2 | 19 | -0.93 | 5.77E-04 | 2.31E-06 |
| ENSG00000213185 | FAM24B | 10 | 0.27 | 6.61E-04 | 2.69E-06 |
| ENSG00000175701 | MTLN | 2 | 0.24 | 1.18E-03 | 5.07E-06 |
| ENSG00000149531 | FRG1BP | 20 | 0.33 | 1.17E-02 | 7.55E-05 |
| ENSG00000015171 | ZMYND11 | 10 | 0.10 | 1.60E-02 | 1.12E-04 |
| ENSG00000140279 | DUOX2 | 15 | -0.91 | 2.38E-02 | 1.92E-04 |
| ENSG00000154760 | SLFN13 | 17 | -0.22 | 3.26E-02 | 2.92E-04 |
| ENSG00000154165 | GPR15 | 3 | 0.33 | 3.35E-02 | 3.15E-04 |
| ENSG00000247516 | MIR4458HG | 5 | 0.22 | 4.86E-02 | 5.74E-04 |

^1 HUGO Gene Nomenclature Committee | 2 Benjamini-Hochberg– adjusted p value^

**TABLE 5|** Differentially expressed genes with adjusted p-value below 0.05 overlapping with top 500 differentially expressed genes in transverse colon in external gene list (Oliva *et al.*, 2020)**.**

| **Ensembl Gene ID** | **HGNC^1^**  **Symbol** | **Chromosome** | **Log2Fold Change** | **Padj^2^** | **pvalue** |
| --- | --- | --- | --- | --- | --- |
| ENSG00000282826 | FRG1CP | 20 | 0.49 | 3.26E-06 | 1.12E-08 |
| ENSG00000240747 | KRBOX1 | 3 | 0.67 | 5.50E-06 | 1.97E-08 |
| ENSG00000022556 | NLRP2 | 19 | -0.93 | 5.77E-04 | 2.31E-06 |
| ENSG00000213185 | FAM24B | 10 | 0.27 | 6.61E-04 | 2.69E-06 |
| ENSG00000175701 | MTLN | 2 | 0.24 | 1.18E-03 | 5.07E-06 |
| ENSG00000149531 | FRG1BP | 20 | 0.33 | 1.17E-02 | 7.55E-05 |
| ENSG00000015171 | ZMYND11 | 10 | 0.10 | 1.60E-02 | 1.12E-04 |
| ENSG00000154760 | SLFN13 | 17 | -0.22 | 3.26E-02 | 2.92E-04 |
| ENSG00000165943 | MOAP1 | 14 | 0.11 | 3.29E-02 | 3.03E-04 |
| ENSG00000104177 | MYEF2 | 15 | 0.17 | 4.06E-02 | 4.22E-04 |
| ENSG00000247516 | MIR4458HG | 5 | 0.22 | 4.86E-02 | 5.74E-04 |

^1 HUGO Gene Nomenclature Committee | 2 Benjamini-Hochberg– adjusted p value^

**FIGURE 2** | Phyper Test (Hypergeometric Distribution in R) comparing 1711 differentially expressed genes in SOCCS to top 500 differentially expressed genes in external gene list (Oliva *et al.*, 2020)**.**

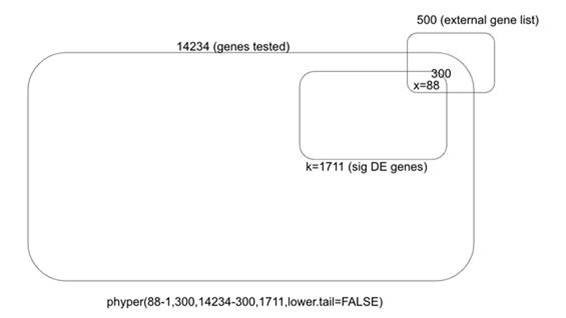
